# Phasor analysis of RGB camera data enables fluorescence microscopy unmixing and brightfield segmentation in a commercial microscope

**DOI:** 10.64898/2026.02.13.705652

**Authors:** Bruno Schuty, María José García, Satya Khuon, Leonel Malacrida

## Abstract

Spectral information plays a crucial role in biological imaging, yet conventional epifluorescence and histological techniques often rely on RGB image acquisition, limiting the resolution of spectrally overlapping components. Here, we present a phasor-based spectral analysis framework adapted for RGB images, enabling unsupervised segmentation and unmixing without the need for hyperspectral systems or sequential acquisition. By applying a discrete Fourier transform to the red, green, and blue intensities at each pixel, we generate a two-dimensional phasor plot where spectral relationships are encoded in modulation and phase. We demonstrate the utility of this approach across three distinct applications: segmentation of lung histology images stained with hematoxylin and eosin to quantify alveolar collapse, analysis of autofluorescence in skin lesions (nevi and melanoma) to highlight pathological spectral signatures, and spectral unmixing in multicolor-labeled U2OS cells to resolve overlapping fluorophores. Our method improves signal separation, reduces noise, and enhances biological interpretability using standard RGB acquisition. These findings establish RGB phasor analysis as a practical and powerful tool for spectral decomposition and segmentation in microscopy, bridging the gap between conventional imaging and advanced spectral analysis.

## Introduction

Microscopy is a cornerstone of biomedical research, enabling visualization of cellular structures, molecular markers, and tissue architecture with high spatial resolution. Over the past decades, advanced imaging modalities such as hyperspectral imaging (HSI), fluorescence lifetime imaging microscopy (FLIM), and multiphoton microscopy have significantly expanded the dimensionality of optical datasets, allowing discrimination of spectrally or temporally overlapping signals [1–3]. Extracting meaningful information from these multidimensional datasets requires robust analytical frameworks capable of simplifying complexity while preserving biological interpretability.

Among such frameworks, the phasor approach has emerged as a powerful model-free method for representing complex optical signals in a two-dimensional Fourier space defined by sine (S) and cosine (G) coordinates [4–6]. In phasor space, similar spectral or temporal behaviors cluster together, and linear combinations of signals follow predictable geometric relationships. This property enables unsupervised classification, spectral unmixing, and quantitative analysis without curve fitting or prior assumptions [2,5,7]. The phasor framework has proven particularly successful in FLIM and hyperspectral microscopy, where it provides intuitive visualization and computational efficiency [1,2,7,8].

Despite its conceptual simplicity, the application of phasor analysis has largely remained restricted to specialized acquisition systems requiring spectral detectors or time-resolved instrumentation [8]. In contrast, RGB imaging is universally available across brightfield microscopy, epifluorescence imaging, histology, and cell biology workflows [9]. However, RGB images are commonly regarded as spectrally limited because they sample emission using only three broad channels, and therefore are typically treated as qualitative representations rather than quantitative spectral datasets [2,10].

Here, we challenge this assumption. We hypothesize that RGB images retain sufficient spectral structure to support phasor-based analysis and that their three-channel intensities can be interpreted as a discrete spectrum suitable for Fourier transformation. By computing the first harmonic phasor coordinates from pixel-wise RGB intensities, each pixel can be mapped into phasor space, where its relative spectral properties are geometrically encoded. Unlike conventional color-based segmentation that relies on intensity thresholds or clustering directly in RGB space, the phasor representation captures inter-channel relationships in a manner that is partially invariant to overall intensity and illumination.

To evaluate the universality and robustness of this approach, we tested RGB phasor analysis across three biologically distinct and increasingly complex scenarios. First, we applied it to brightfield histological images of lung tissue from a murine model of acute lung injury (ALI), where quantification of alveolar collapse and tissue remodeling is traditionally performed using stereological methods [11–17] or intensity-based segmentation approaches [17–20]. Second, we analyzed autofluorescence images of melanocytic lesions, where endogenous fluorophores such as NADH, FAD, collagen, elastin, and melanin generate subtle spectral variations associated with metabolic and structural heterogeneity [21–25]. Third, we implemented RGB phasor spectral unmixing in multicolor epifluorescence microscopy, a setting where overlapping fluorophore emission and channel crosstalk typically require hyperspectral acquisition or matrix-based unmixing strategies [26–29].

By spanning brightfield, label-free fluorescence, and multicolor fluorescence imaging, these applications provide a comprehensive assessment of whether RGB phasor analysis can bridge conventional microscopy and advanced spectral methodologies. Together, they establish RGB phasor analysis as an accessible extension of the phasor framework, capable of transforming standard color images into quantitative spectral representations without specialized hardware.

## Materials and Methods

### Materials

Tissue samples for autofluorescence imaging were obtained from patients diagnosed with melanoma or benign nevi. Paraffin-embedded sections were prepared at 5 µm thickness and mounted on standard microscope slides using Canada Balsam (Biopack) as a mounting medium to ensure refractive index matching. The studies involving humans were approved by Prof. Dr. Raúl Ruggia, Coordinator of the Research Ethical Committee at the Hospital de Clinicas, Universidad de la Republica, Uruguay. The studies were conducted in accordance with the local legislation and institutional requirements. The participants provided their written informed consent to participate in this study.

Mouse lung tissue was collected from two experimental groups: control mice under a normal diet (B6 ND) and mice treated with intratracheal lipopolysaccharide (LPS) under a normal diet (B6 ND + LPS). The lungs were fixed in 10% neutral buffered formalin and embedded in paraffin. Serial histological sections were stained with hematoxylin and eosin. The animal work was evaluated and approved by the National Ethics Committee with the animal protocol for Animal Use (Nº 070153-000571-19).

U2OS human osteosarcoma cells were stained using DAPI (nuclear label), rabbit anti-nuclear Lamin A with goat anti-rabbit Alexa Fluor 555 (nuclear lamina) and mouse anti-Tubulin cl DM1A and with goat anti-mouse Alexa Fluor 488 (cytoskeleton). All samples were prepared on standard glass coverslips.

For RGB-phasor acquisition, we used an Olympus IX81 widefield epifluorescence microscope, with simple modifications. We installed a Kiralux 5.0 MP Color CMOS Camera (CS505CU). The camera was installed in the third port of the microscope at the eyepiece. For imaging the RGB-camera was run at 8-bit depth. We modified the regular Olympus excitation-emission filters by removing the emission filter from the UV-MNU2 filter to allow the collection of the entire emission spectrum. While for cell imaging, a commercial RGB filter cube for excitation and emission was used. The cube 69015 ET – DAPI/Green/Red FISH consisted of a multi-bandpass filter in excitation (69015x) and emission (69015m), and a dichroic mirror (69015bs) from Chroma Technology Corp.

Acquisition was controlled using the ThorImage®CAM Software (v3.7.0). We used a UPlanFLN 10x NA 0.30 FN26.5 objective and an Olympus LUCPlanFLN 40x NA 0.60 FB22 oil immersion objective for melanoma and lung tissue, respectively. For RGB demixing experiments, an Olympus PlanApo N 60x objective with NA 1.42, oil immersion 0.17/FN26.5 USI 2 BFP1 was used. For all experiments, the camera was set with a 100 ms exposure time, a frame rate of 53.195 FPS, and images were generated from 4-frame averaging. Color gains of 2.63, 1.0, and 3.41 were used for red, green, and blue channels, respectively.

A Zeiss LSM 880 confocal microscope with a 10x Plan-APOCHROMAT NA 0.45 and a spectral detector, using 405 nm laser excitation, was employed for label-free autofluorescence imaging. All image processing and quantitative analysis were performed using Python 3.11. The main libraries used were PhasorPy [30], NumPy [31], SciPy [32], and Scikit-Image [33]. Simulations were also performed in this environment.

### Simulations

A synthetic RGB color wheel (256×256 pixels) was generated based on the HSV color model. Hue values were computed from the angular coordinates of each pixel, saturation was derived from radial distance (clipped to 1), and value was set to 1 for full brightness. The HSV image was converted to RGB and masked outside the unit circle to produce a white background.

Phasor coordinates were calculated using the *phasor_from_signal* function from the *PhasorPy* library (v0.6) [30]. The phasor representation of the image was used to segment regions via manual cursor selection and clustering. Unsupervised segmentation was evaluated using *KMeans* and *GaussianMixture* models from *Scikit-learn*, with different covariance settings (full, tied, diag, spherical).

Spectral unmixing from [29] was applied to decompose the image into red, green, and blue components, using their known pure coordinates in RGB space: blue = [1, 0, 0], green = [0, 1, 0], and red = [0, 0, 1]. The unmixed phasor data was used to reconstruct the original RGB image, validating the phasor-based decomposition under ideal conditions.

### Sample preparation and Imaging

#### Hematoxilin Eosin

C57BL/6 male mice (8 weeks old) maintained on a normal diet were divided into two experimental groups: (1) control without treatment (B6 ND) and (2) mice receiving an intratracheal instillation of lipopolysaccharide (LPS) (B6 ND + LPS). LPS from *Escherichia coli* O55:B5 (Sigma-Aldrich, St. Louis, MO) was administered at a dose of 3 µg/g body weight under isoflurane anesthesia, following the protocol of [34]. Control animals received an equivalent volume of sterile saline. Forty-eight hours post-instillation, lungs were collected and fixed in 10% neutral buffered formalin. Fixed tissues were then dehydrated through a graded ethanol series (50%, 70%, 96%, and isopropanol; 30 min each), followed by a 30-minute step in chloroform, and subsequently embedded in paraffin. Sections were stained with hematoxylin and eosin (H&E) according to standard histological protocols. H&E sections were imaged using an Olympus IX81 epifluorescence microscope operated in transmission mode, equipped with an Olympus UPlanFL N 40×/0.75 NA objective. Images were acquired with a Thorlabs Kiralux 8-bit RGB camera controlled by the manufacturer’s software. The microscope was fitted with an ASI motorized XY stage (model MS-2000). For each lung sample, ten random regions were imaged under identical acquisition settings. In total, 100 RGB images were analyzed per study group.

#### Label-free

As previously described in [27] Formalin-fixed, paraffin-embedded (FFPE) sections of nevi and melanomas were cut at 5 μm thickness. The sections were mounted on glass slides and incubated at 60 °C for 30 minutes to ensure proper adhesion. To preserves optical properties and enhance imaging quality, a drop of Canada Balsam (Biopack) was applied before placing a coverslip, providing refractive index matching for label-free autofluorescence microscopy. Images were acquired with the Olympus IX81 widefield epifluorescence microscope was configure using the UV-MNU2 filter without the emission filter. The emission filter was removed to collect the full emitted fluorescence spectrum of the sample. As a result, the RGB image captured by the color camera represented a broadband integration of the emission light, effectively compressing the spectral information into three channels. Images were saved in TIFF format at 8-bit resolution. To obtain reference spectral data from the same sample, hyperspectral images were acquired using a Zeiss LSM 880 confocal microscope equipped with a Gallium-Arsenide Phosphide (GaAsP) photomultiplier tube (PMT). The detector was set with a gain between 520 V and 550 V and collected emission spectra from 423 nm to 723 nm in 30 channels, with a spectral resolution of 10 nm. Excitation was performed using a 405 nm laser, and a 405 dichroic mirror was placed in the optical path to reflect the excitation light.

#### Fluorescence imaging

U2OS human osteosarcoma cells were stained with three fluorescent markers: DAPI for nuclear DNA, mouse monoclonal Anti-TUBA4A (TUBA1) Antibody (Merck, Cat # T9026), and counterstained with goat anti-mouse IgG Alexa Fluor™ 488 (Thermo Fisher Scientific, Cat #A28175) for tubulin, and rabbit monoclonal Anti-Lamin A/C Antibody, clone 2B10 ZooMAb® (Merck, Cat # ZRB1054) and goat anti-rabbit IgG Alexa Fluor™ 555 for the nuclear envelope. Two types of samples were prepared: multicolor-labeled cultures containing all three fluorophores, and single-labeled control cultures used as reference components for spectral unmixing. The Olympus IX81 inverted epifluorescence microscope was configured with the cube 69015 ET – DAPI/Green/Red FISH to excite and collect fluorescence from our three labels (DAPI, Tubulin Alexa Fluor 488, and Lamin A Alexa Fluor™ 555). We used the ThorLabs Kiralux RGB camera for image capture. An Olympus PlanApo N 60x objective with NA 1.42 was used. All images were acquired under identical exposure conditions and stored as TIFF format and 8-bit resolution. The acquisition was performed using ThorImage^®^CAM Software, and no spectral separation beyond the RGB color channels was applied.

### Data analysis

#### Phasor Analysis in RGB imaging

The phasor transform provides a compact, model-free representation of a signal by mapping it into a two-dimensional phasor space defined by its real (G) and imaginary (S) components [2, 5]. For a discrete signal with N intensity values *I*_*k*_, the first harmonic phasor coordinates are computed as shown in Equations (1) and (2) where, N is the number of spectral channels or time bins in the signal, k is the index of the channel and *I*_*k*_ signal intensity at channel k.

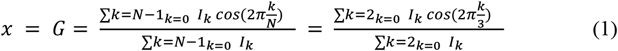

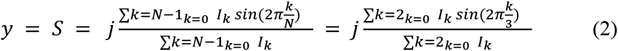

The resulting phasor (G, S) lies within the unit circle and encodes spectral shape: the angle reflects the spectral center (or phase), and the distance to the origin (modulation) relates to spectral width or purity. In this work, we apply the phasor transform with N=3, corresponding to the red, green, and blue (RGB) channels of standard color cameras.

#### Brightfield images

were analyzed using phasor-based segmentation to differentiate morphologically and colorimetrically distinct components. The phasor transform was applied to each RGB image. The distribution of these coordinates captures the spectral composition of the tissue, with nuclei (blue-violet), red blood cells (reddish), extracellular tissue (pinkish), and background (white/alveolar space) occupying distinct regions. To segment these components, K-means clustering was applied to the phasor coordinates with k = 4 as described by [18]. This algorithm was selected due to its computational efficiency, making it suitable for processing large image batches. The resulting cluster labels were mapped back onto the spatial domain to reconstruct segmented images, where each pixel was assigned to one of the four identified chromatic classes. Following segmentation, the alveolar area was quantified. The cluster corresponding to the background (alveolar space) was identified based on its phasor coordinates and spatial distribution, and its pixel count was compared to the total tissue area to compute the alveolar space percentage for each image.

For each animal (n = 10 images per mouse, 10 mice per group), the mean alveolar space was calculated. Two experimental groups were compared: B6 ND and B6 ND+LPS. The results were visualized using boxplots (one value per mouse) and violin plots to illustrate the distribution within each group. To evaluate statistical differences, the distribution of alveolar scores in each group was assessed for normality by computing the coefficient of skewness and applying the Shapiro-Wilk test. If the data met the assumptions of normality, an independent two-sample t-test was performed to compare the means of both groups, and the resulting p-value (p) was reported.

To assess the discriminative potential of the alveolar space metric, a Receiver Operating Characteristic (ROC) curve was generated. The Youden Index (J = Sensitivity + Specificity - 1) was used to determine the optimal threshold for classification between B6 ND and B6 ND+LPS groups. A simple logistic regression model was then trained using the alveolar space percentage as a single input feature. The model’s performance was evaluated using a confusion matrix, accuracy, sensitivity, and specificity metrics on the full dataset.

#### Autofluorescence images

For each image, phasor coordinates were computed by applying the Fourier transform to each pixel’s spectral profile. In the case of RGB images, the 3-channel vector [R, G, B] was interpreted as a discrete spectrum and transformed into a phasor representation using the first harmonic as described by Equations 1 and 2. A graphical phasor plot was generated for each image, and cursors were manually placed to define regions of interest based on prior knowledge of fluorophore distributions. Each pixel was assigned to a pseudocolor based on its location within the phasor plot, and a pseudocolor image was reconstructed by mapping these classifications back to the spatial domain.

#### Fluorescence images

were first preprocessed using an intensity thresholding step. A channel-wise histogram analysis was performed to define specific intensity ranges for each color channel, effectively removing pixels with very low signal and those with saturated values in any channel. Following preprocessing, phasor coordinates were calculated for each pixel using Equations 1 and 2, which transform the RGB intensity values into phasor space for spectral decomposition. Spectral unmixing was then performed using the RGB-phasor unmixing algorithm [30]. The algorithm computes unmixed fraction images for each channel, based on reference phasor positions derived from the center of mass of pure component images. To determine these references, Otsu thresholding was applied to the average intensity image of each pure fluorophore, and the resulting mask was used to extract the phasor pixels corresponding to each component. Photon count images were subsequently generated by modulating the original intensity of each pixel by the unmixed fractional values of the corresponding channel. For a three components sample the Linear System is given by Equation 3, where (*G*_*i*_, *S*_*i*_) and the center of mass of the phasor distribution of for each pure component *i* (1, 2, 3) and *G*_*exp*_ and *S*_*exp*_ are the phasor coordinates for each measure spectrum.

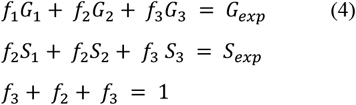

This approach improves the estimation of fluorophore-specific signals by incorporating both spectral and intensity information. An RGB image was reconstructed using the unmixed and intensity-modulated channels. A region of interest (ROI) was manually selected from an area containing overlapping signals from all three fluorophores. Intensity profiles were then extracted along a defined line within the ROI, allowing comparison of the signal distribution before and after unmixing.

## Results

### Simulations

Starting from a simulated RGB color space, we applied phasor analysis to explore its utility under spectrally limited conditions with only three channels. An 8-bit RGB image was transformed pixel-wise to generate a phasor plot (Fig. 1.C), where primary (R, G, B) and secondary colors (C, M, Y) were coherently distributed. We then tested three classical phasor methods. First, manual cursors selected primary colors (Fig. 1.D). Second, a Gaussian mixture model clustered the phasor plot into three spectral regions automatically (Fig. 1.E). Third, spectral unmixing modeled a phasor point as a linear combination of pure components, estimating their contributions (Fig. 1.F). The resulting images (Figs. 1.G–I) show consistent segmentation across the three approaches. Overall, this simulation demonstrates that even with RGB data, phasor analysis enables effective decomposition, visualization, and segmentation of chromatic components.

**Fig 1.**
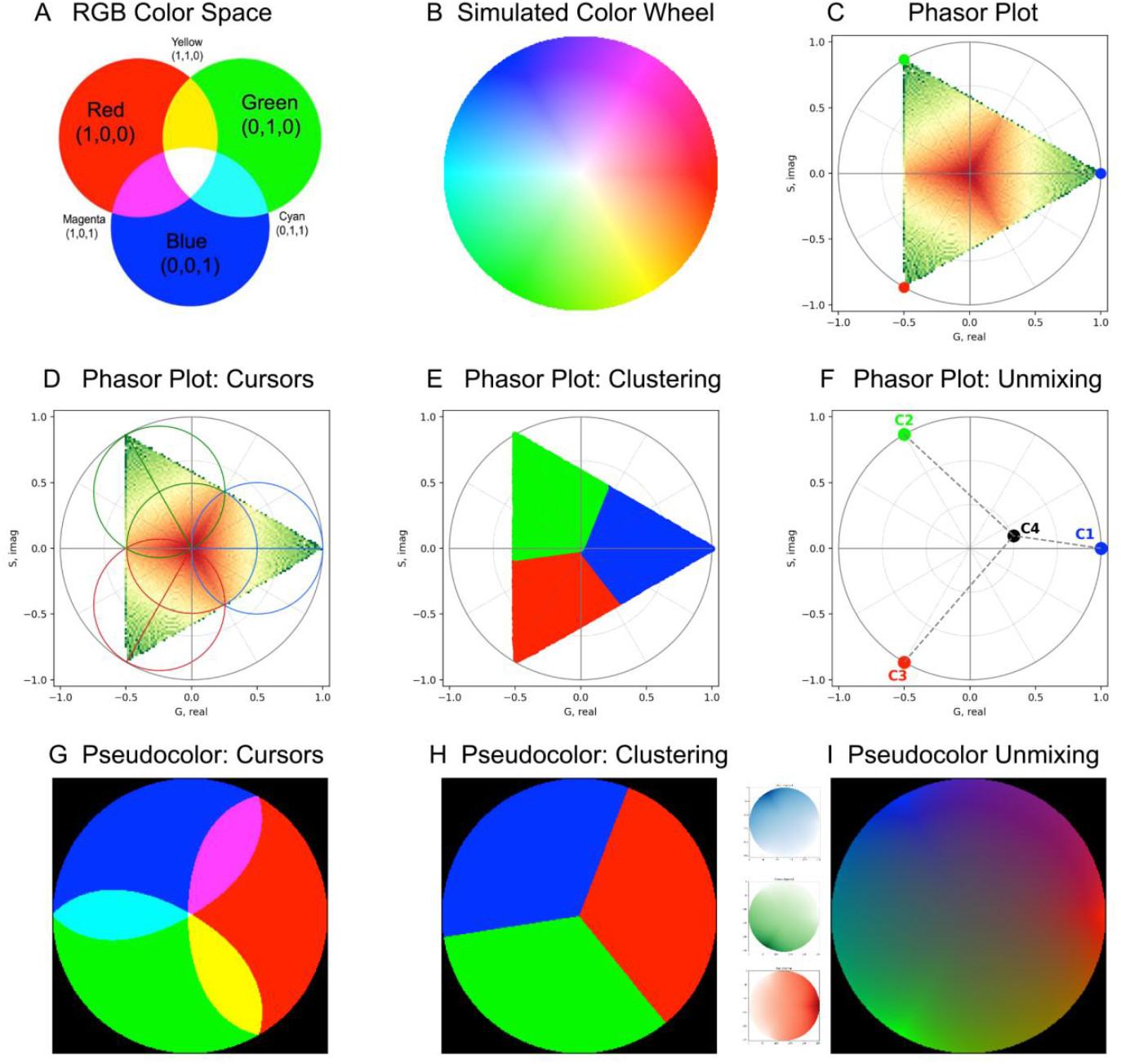
Phasor analysis for an RGB color space. A) RGB color space, where Red, Green and Blue are the base of color compositions, those are digitally represented by (1,0,0), (0,1,0) and (0,0,1) respectively. Cyan, Magenta and Yellow are combinations of those which are represented by (0,1,1), (1,0,1) and (1,1,0), the presence of the three components generates the white color (1,1,1). B) Color Wheel of an RGB space simulated with 8 bits depth. C) Phasor plot obtained from B) with the pure components (RGB) represented on it and the combinations of them (CMY). D) Phasor plot with three cursors to select regions of interest in the phasor plot obtained in C). E) Clusterized Phasor plot of C) with a Gaussian mixture model, implemented with the Scikit-Learn Library. F) Representations of the spectral unmixing principle, given a component C4, C4 is a linear combination of the pure components (C1, C2, C3) relying on the image. G) Pseudocolor image obtained with cursors, intersection areas are represented with the combination of pure color components as shown in A). H) Pseudocolor image obtained from clusterized phasor plot. I) Image obtained for pure unmixed components Blue, Green and Red and the color image obtained unmixed from B) with the spectral unmixing method.

**Fig 2.**
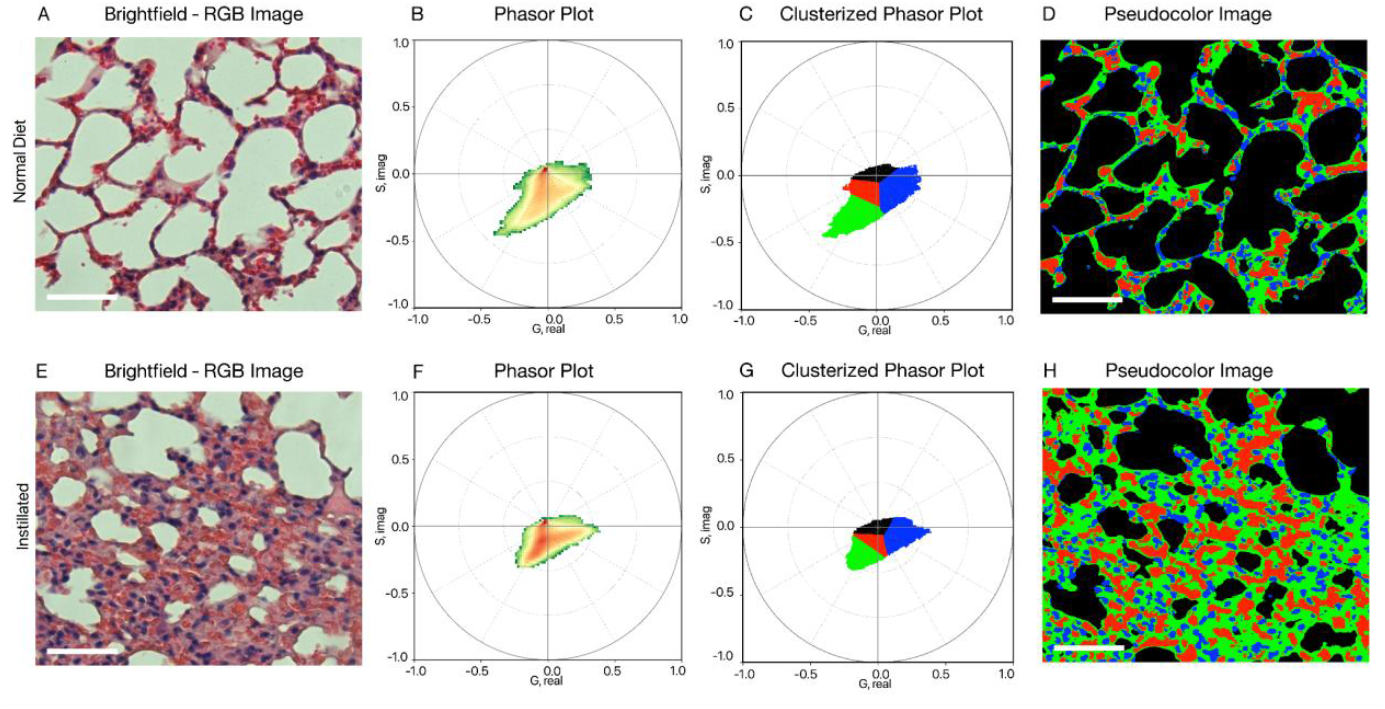
RGB Phasor Analysis of alveolar architecture in mouse lung tissue. (A, E) Representative RGB images of H&E-stained lung sections from normal diet (ND) and instilled mice. (B, F) Phasor plots derived from the RGB images showing the spectral distribution of pixel intensities. (C, G) Clustered phasor plots highlighting four spectral classes (red, blue, green, black) used for segmentation. (D, H) Segmented RGB images using phasor-based clustering to identify alveolar spaces (black) and tissue components. A 50 μm scale bar is shown.

### Brightfield

Representative RGB images of lung tissue from mice B6 ND and B6 ND+LPS conditions are shown in panels A and E. Corresponding phasor plots (panels B and F) and their clustered versions (panels C and G) highlight the spectral separation of tissue components, including hematoxylin-stained nuclei, eosin-stained fibrous tissue, red blood cells, and alveolar spaces. These spectral classes enabled segmentation of alveolar structures, shown as masks overlaid on the original RGB images (panels D and H). Morphometric quantification of alveolar area across 100 images per group is summarized using grouped boxplots (Fig 3.A) and violin plots (Fig3.B). Both B6 ND and B6 ND instilled group distributions passed the Shapiro-Wilk test for normality (W = 0.929 and 0.507, respectively), allowing the use of parametric testing. A two-tailed unpaired t-test revealed a highly significant difference in alveolar space area between groups (p < 0.0001), with a large effect size (Cohen’s d = −3.61). Classification based on alveolar area was performed using logistic regression, with excellent performance indicated by the ROC curve (AUC = 0.993, optimal threshold = 0.596, Fig 3.C) and confusion matrix (Fig 3.D), achieving 97% accuracy and an F1 score of 0.97. These results demonstrate that RGB-based phasor segmentation can reliably extract morphometric features with diagnostic potential, later discussed in this work.

**Fig 3.**
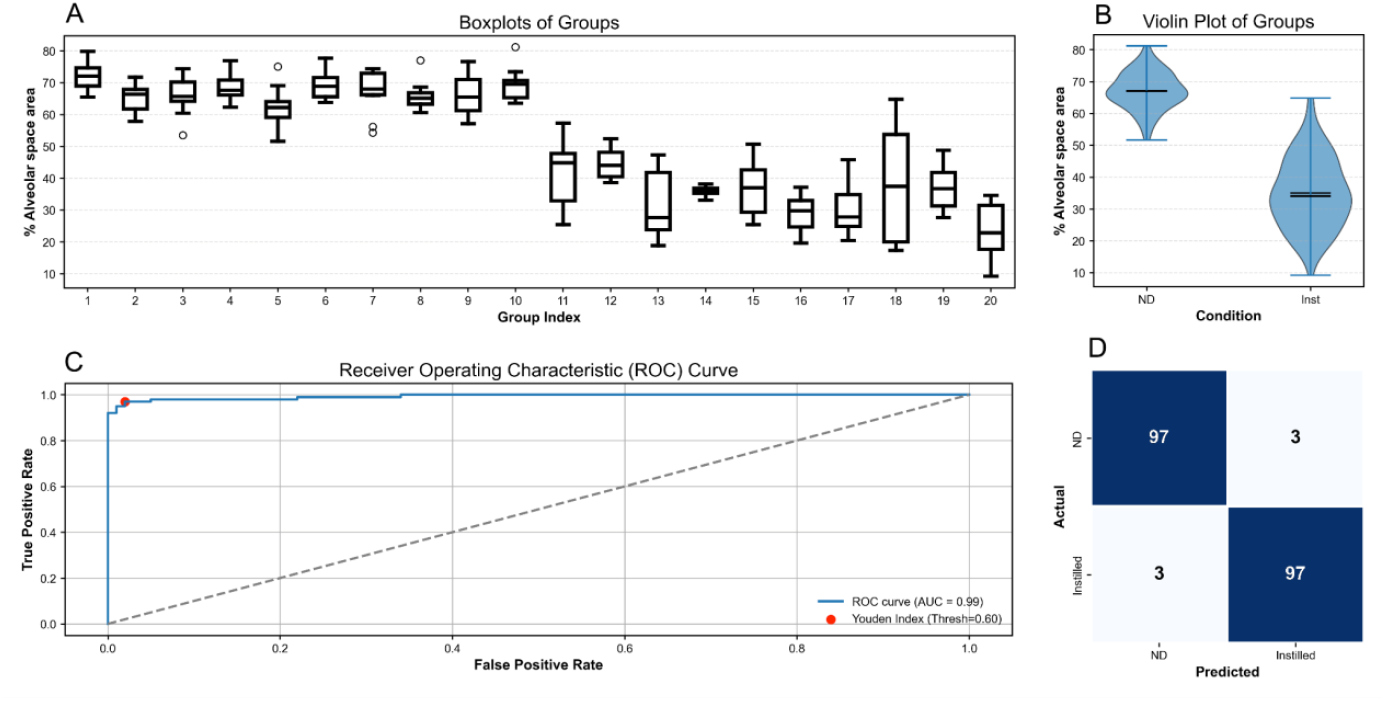
(A) Boxplots showing the percentage of alveolar space area for each of 20 individual mice (10 ND, 10 instilled and 10 images per mice). (B) Violin plots comparing the alveolar space area between ND and instilled groups. (C) Receiver operating characteristic (ROC) curve of the logistic regression model distinguishing ND vs. instilled lungs based on phasor-derived features (AUC = 0.99). (D) Confusion matrix showing the classification performance of the same model (97% accuracy for both ND and instilled groups).

### Autofluorescence

To assess the performance of phasor-based analysis on autofluorescence data acquired through RGB and hyperspectral imaging (HSI), we analyzed matched tissue samples of nevi and melanoma. Figure 4 presents representative examples from each class. Panels A-D show the original autofluorescence images in both RGB and HSI modalities. Their corresponding phasor plots (E-H) reveal the spectral pixel distributions in phasor space. Cursor-based segmentation was applied to identify major spectral domains, and the resulting pseudocolor maps (I-L) highlight spectrally distinct regions within the tissue. In addition to these qualitative visualizations, we applied principal component analysis (PCA) to the phasor coordinates, extracting parameters such as elongation, variance, and ellipse area. The results showed higher elongation and dispersion in melanoma compared to nevi, with moderate differences between RGB and HSI modalities. These quantitative comparisons are detailed in the Supplementary Material.

**Fig 4.**
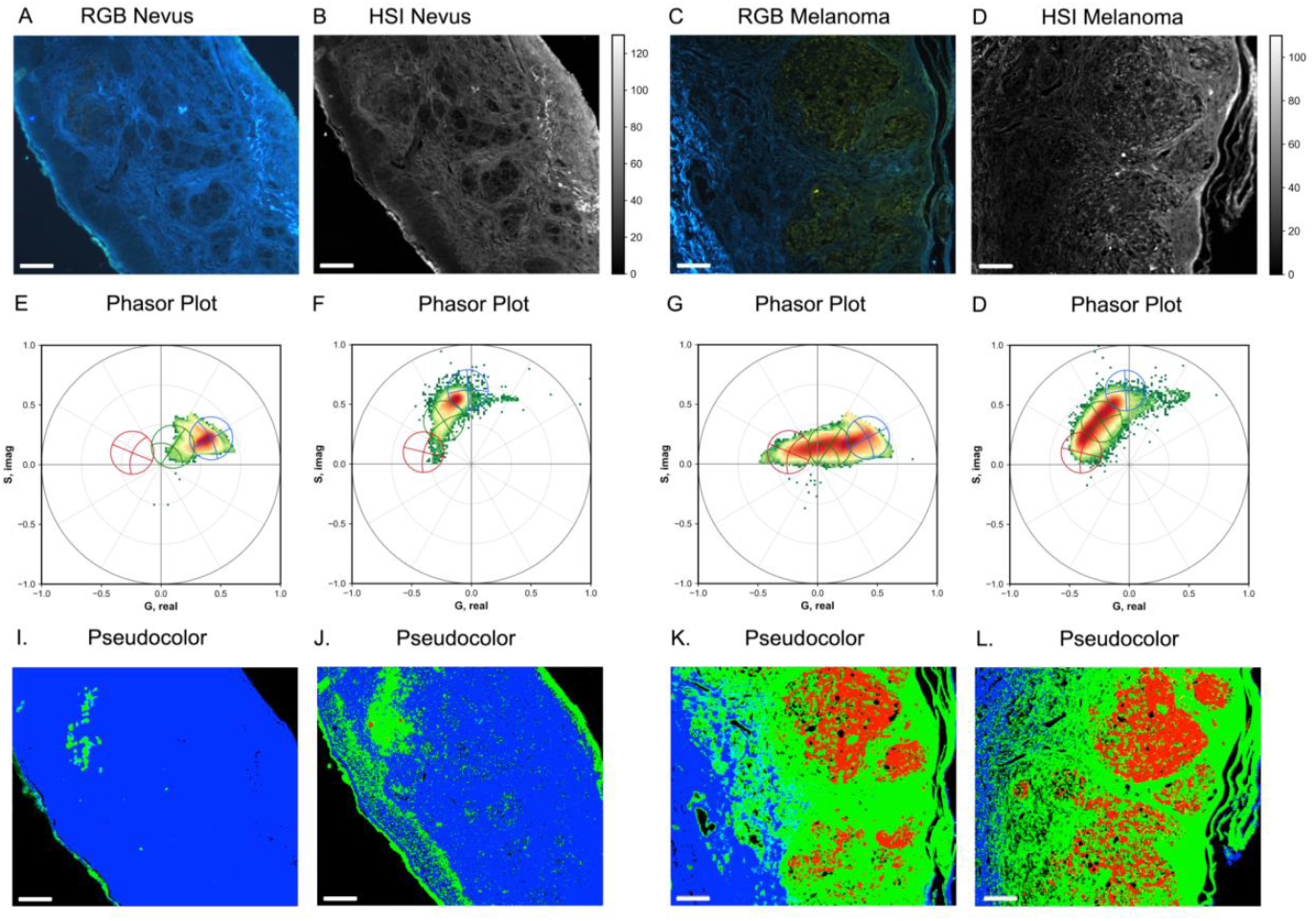
Phasor analysis of autofluorescence in nevi and melanoma using RGB and HSI images. (A–D) RGB and HSI autofluorescence images of nevus and melanoma tissue, respectively. (E–H) Corresponding phasor plots for each image, showing the spectral distribution of pixels in phasor space. (I–L) Pseudocolor segmentations obtained from phasor cursor selection, illustrating distinct spectral regions associated with tissue features. RGB and HSI modalities were analyzed for the same tissue regions to compare segmentation performance and phasor behavior. The scale bar represents 100 μm.

### RGB Spectral Unmixing

To demonstrate the capabilities of the RGB phasor unmixing approach, we applied it to a multicolor epifluorescence image of fixed cells. Figure 5 shows the initial steps of the analysis. The labels and molecular markers were chosen to stress the overlapping signal from the different components. For instance, DAPI and Laminin Alexa Fluor™ 555 spatial overlaps and a strong bleeding from DAPI into Green and Red channel can be identified at Figure 5 B, C and D. Separation of the blue, green, and red channels (Fig. 5B–D) reveals the contributions of each color, while the histograms (Fig. 5E) highlight their intensity distributions. The phasor plot of the image (Fig. 5F) illustrates how pixels are distributed in phasor space, where cursors were placed to define spectral regions. Based on these selections, a pseudocolor representation was generated (Fig. 5G), providing a first visualization of distinct spectral contributions.

**Fig 5.**
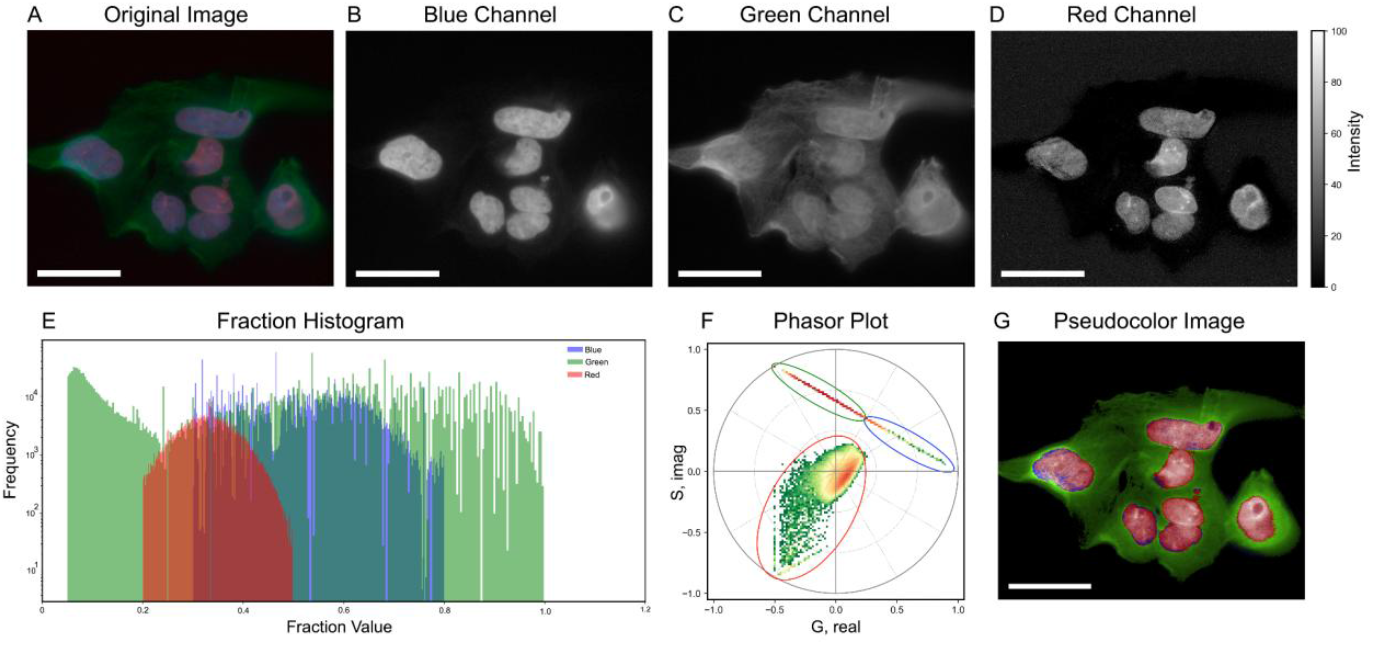
(A) Original RGB image of the sample. (B–D) Individual blue, green, and red channels extracted from the RGB image. (E) Histograms of pixel intensity distributions for each channel are shown in (B–D). (F) Phasor plot of the RGB image, illustrating the distribution of pixels and the selected cursor positions used to define spectral regions. (G) Pseudocolor representation generated by assigning pixel clusters from the phasor plot in (F) to their corresponding regions in the image. The scale bar represents 50 μm.

The subsequent steps are summarized in Figure 6. The unmixed fractions for the blue, green, and red components (Fig. 6A–C) show clear spatial patterns, which are also evident in the intensity maps of each unmixed channel (Fig. 6D–F). When compared with the original RGB image (Fig. 6G), the reconstructed unmixed RGB image (Fig. 6H) reveals a noticeable reduction in signal overlap. Finally, the intensity profiles along a selected region (Fig. 6I) illustrate how the separation achieved by the phasor analysis reduces crosstalk between channels and enhances the visualization of individual components.

**Fig 6.**
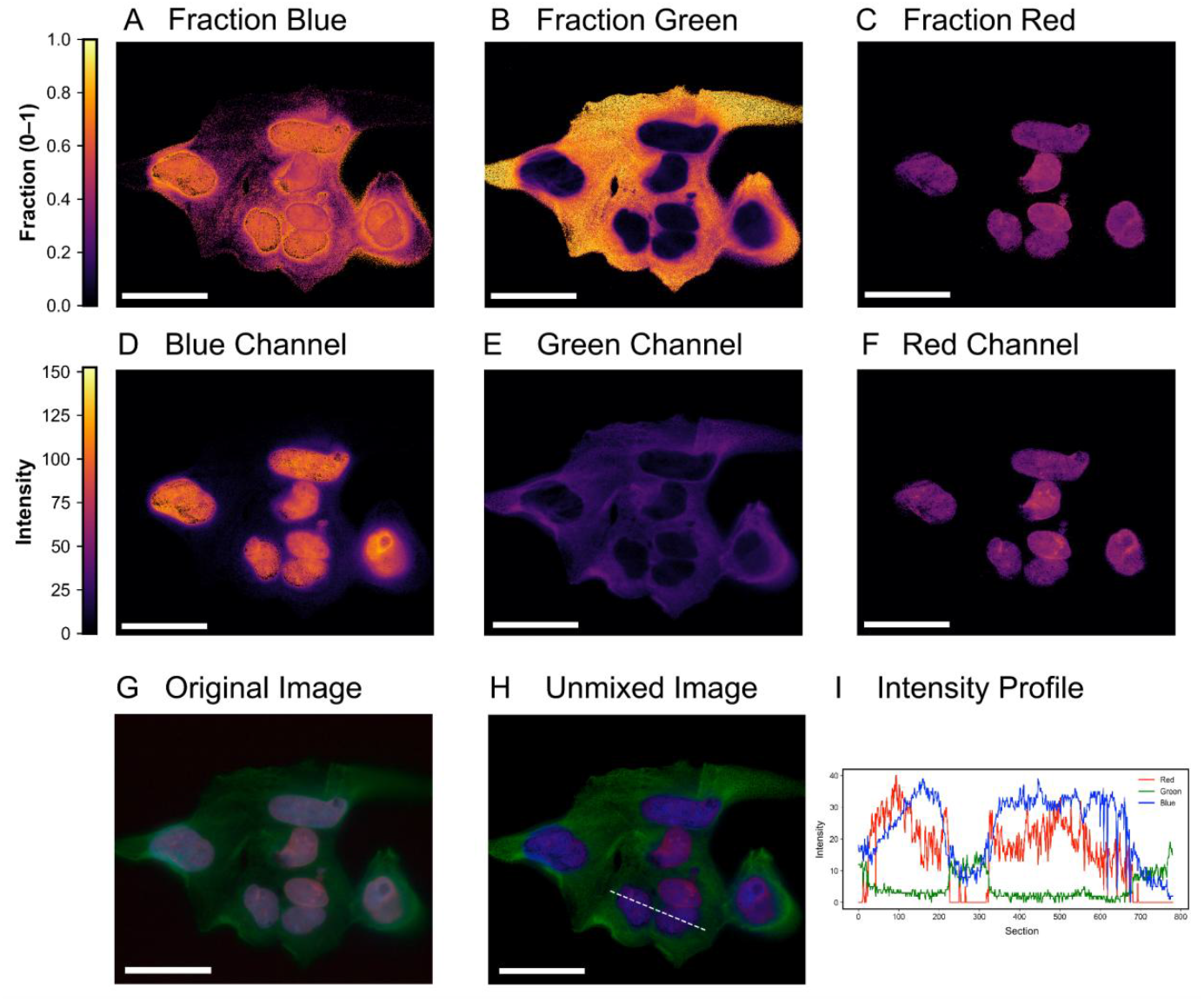
(A–C) Unmixed fractions for the blue, green, and red components. (D–F) Intensity maps of the unmixed blue, green, and red channels. (G) Original RGB image. (H) Reconstructed unmixed RGB image with a dashed white line indicating the region selected for intensity profiling. (I) Intensity profiles of the unmixed RGB channels along the line drawn in (H), showing the spatial distribution and relative overlap of the components. The scale bar represents 50 um.

## Discussion

The central finding of this work is that three-channel RGB data preserve sufficient spectral geometry to enable meaningful phasor analysis. Although RGB sampling is coarse compared to hyperspectral acquisition, the linear-combination properties that define the phasor space remain valid [4–6]. Across simulations and biological applications, we observed that primary and mixed color components occupy predictable positions in phasor space, and that clustering, segmentation, and unmixing follow the same geometric rules described for lifetime and hyperspectral phasor methods [1,2,5,7,8]. These results support the view that phasor analysis is fundamentally a geometric transformation of relative spectral relationships rather than a technique dependent on high spectral resolution.

Our simulations provide theoretical grounding for this interpretation. The triangular distribution of primary and secondary colors in the RGB wheel reproduces the linear mixing behavior previously described in phasor theory [4,6], confirming that coarse spectral sampling does not eliminate geometric structure. Instead, RGB data serve as a compressed spectral representation that preserve essential relationships. This finding extends previous demonstrations of spectral phasor robustness [2,7] and reinforces the method’s conceptual generality.

Recent work has demonstrated the applicability of phasor-based analysis in digital pathology, particularly for the quantitative assessment of iron overload in Perls’ Prussian Blue-stained liver biopsies, where spectral and spatial autocorrelation phasor representations were combined to extract concentration and granule size information from RGB whole-slide images [35]. That study highlights the strength of phasor geometry for explainable, unsupervised segmentation in a clinically relevant context.

While that approach focused on a specific staining system and pathology-driven quantification, our work expands the conceptual scope of RGB phasor analysis beyond single-disease applications. By establishing a simulation-based theoretical foundation and demonstrating applicability across brightfield histology, label-free autofluorescence imaging, and multicolor fluorescence unmixing, we position RGB phasor analysis as a modality-independent geometric framework rather than a task-specific algorithm. In this sense, the present study generalizes and extends the utility of RGB phasor analysis beyond targeted digital pathology applications to a broader range of biological imaging contexts.

In brightfield histology, RGB phasor analysis enabled unsupervised segmentation of lung architecture with high reproducibility. Traditional stereological approaches remain the gold standard for morphometric assessment [11,15–17], yet they are labor-intensive and subject to operator variability. Intensity-based thresholding and clustering methods improve automation but remain sensitive to staining and illumination variations [18–20]. In contrast, phasor coordinates reflect relative channel contributions rather than absolute intensities, producing reproducible clustering of nuclei, erythrocytes, fibrous tissue, and alveolar spaces. The significant reduction in alveolar fraction in LPS-treated lungs is consistent with established models of acute lung injury [12–14], and the high discriminative performance (AUC = 0.99) indicates that phasor-derived morphometric features may provide a robust alternative to manual quantification. In line with prior efforts to enhance the analytical utility of RGB histology [9,10], our results demonstrate that phasor transformation expands the quantitative potential of standard brightfield imaging without altering acquisition protocols.

In autofluorescence imaging of melanocytic lesions, RGB phasor distributions mirrored the spectral heterogeneity observed with hyperspectral imaging. Spectral phasor analysis has previously revealed metabolic and structural differences driven by endogenous fluorophores such as NADH, FAD, collagen, and melanin [21–25]. Consistent with these reports, melanoma samples exhibited more elongated and dispersed phasor distributions than nevi, reflecting increased spectral heterogeneity. Although RGB data cannot resolve fine spectral peaks with the precision of HSI [3,24,25], global descriptors such as elongation, variance, and entropy retained biological relevance. This suggests that diagnostic information may be encoded in the geometry of spectral relationships rather than in high-resolution spectral detail alone. Therefore, RGB phasor analysis offers a pragmatic alternative in settings where hyperspectral instrumentation is unavailable or impractical.

In multicolor epifluorescence microscopy, the RGB phasor unmixing strategy exploited the linear mixing behavior intrinsic to phasor space [4–6,29]. Spectral overlap and crosstalk are well-known limitations of conventional fluorescence microscopy [26–28], typically addressed through sequential acquisition or matrix-based algorithms. By projecting pixel phasor coordinates onto reference vectors derived from pure fluorophores [29], we achieved separation of overlapping signals from single-exposure RGB images. While quantitative precision is constrained by the limited number of spectral channels, the reduction in crosstalk and improved spatial contrast demonstrate practical utility. This approach may be particularly valuable in high-throughput, live-cell, or resource-limited environments where acquisition simplicity is essential.

Several limitations should be acknowledged. The spectral resolution of RGB data inherently restricts fine biochemical discrimination, and variations in illumination spectrum, camera response, or staining protocols may influence phasor coordinates. As described for lifetime and multispectral phasor analyses [5,7,10], calibration strategies and internal normalization may enhance reproducibility across systems. Future developments could integrate machine-learning classifiers trained directly in phasor space, combining geometric interpretability with scalability.

Overall, this study demonstrates that RGB phasor analysis preserve the analytical strength and interpretability of established phasor methodologies [1–8] while substantially lowering the technical barrier to adoption. By revealing that conventional RGB images contain recoverable spectral geometry, we redefine color microscopy as a compressed spectroscopic modality rather than a purely qualitative representation. This conceptual shift broadens access to phasor-based analysis and supports its integration into routine biomedical imaging workflows.

## Conclusion

This work establishes RGB phasor analysis as an accessible and conceptually unified extension of the phasor framework, demonstrating that three-channel color images retain sufficient spectral geometry for quantitative interpretation. Across brightfield histology, autofluorescence imaging, and multicolor fluorescence microscopy, we show that the linear properties of phasor space remain valid even under coarse spectral sampling. Rather than viewing RGB acquisition as a limitation, our results reinterpret it as a compressed spectral encoding that can be mathematically expanded through Fourier transformation.

By bridging simulations with diverse biological applications, we demonstrate that phasor analysis functions as a modality-independent geometric language for optical data. The ability to perform segmentation, pseudo-spectroscopy, and spectral unmixing directly from RGB images transforms conventional color microscopy into a quantitative analytical tool without requiring specialized hardware. This substantially lowers the technical barrier to advanced spectral analysis and broadens accessibility beyond laboratories equipped with hyperspectral or time-resolved instrumentation.

Importantly, RGB phasor analysis does not aim to replace high-resolution spectral systems. Instead, it offers a pragmatic alternative when acquisition complexity, cost, or throughput constraints limit the deployment of hyperspectral technologies. The framework preserve interpretability while enabling automation, making it compatible with open-source ecosystems such as PhasorPy [30] and modern computational pipelines.

Looking forward, several avenues can further expand the impact of this approach. Standardization strategies for illumination and camera calibration may improve inter-system reproducibility. Integration with machine-learning models trained directly in phasor space could enhance classification performance while retaining geometric interpretability. Additionally, the development of optimized RGB or narrowband filter sets may increase effective spectral resolution without sacrificing accessibility.

Ultimately, RGB phasor analysis reframes conventional color microscopy as a spectroscopically informative modality. By revealing the recoverable spectral structure embedded in everyday RGB images, this work extends the reach of phasor methodology from advanced biophotonics laboratories to routine biomedical, clinical, and educational settings, supporting more inclusive and quantitative imaging practices.

## Acknowledgements

The authors declare no conflict of interest. We would like to thank Ana Laura Suárez, technician in anatomo-pathology at the Advanced Bioimaging Unit, for preparing the hematoxylin and eosin-stained lung tissue samples from mice. We also thank Verónica Lezue for her assistance in preparing the label-free melanoma and nevus tissue samples. We are grateful to the members of the Advanced Bioimaging Unit for their insightful discussions and contributions throughout this project. Finally, we thank the Dermatology Division of the Hospital de Clínicas for kindly providing the clinical tissue samples used in this study. LM is a researcher at PEDECIBA (Programa de Desarrollo de las Ciencias Básicas, Uruguay) and the Sistema Nacional de Investigadores (SNI-ANII, Uruguay). LM is supported by grants #2020–225439, #2021– 240122, and #2022–252604 from the Chan Zuckerberg Initiative DAF, an advised fund of the Silicon Valley Community Foundation.

