## Supplementary Material for "Phasor analysis of RGB camera data enables fluorescence microscopy unmixing and brightfield segmentation in a commercial microscope"

### Phasor Analysis of RGB Imaging Enables Unmixing and Tissue Segmentation Across Fluorescence and Brightfield Microscopy: Supplementary Material

Bruno Schuty [1, 2], María José García [1, 3], Satya Khuon[4], Leonel Malacrida [1,3*]

[1] Advanced Bioimaging Unit, Institut Pasteur Montevideo and Universidad de la República, Montevideo, Uruguay.

[2] Present address: Department of Biomedical Engineering, University of California, Irvine, Irvine, CA, USA

[3] Unidad Académica de Fisiopatología, Hospital de Clínicas, Facultad de Medicina, Universidad de la República, Montevideo, Uruguay.

[4] Advanced Imaging Center and Integrative Imaging, Howard Hughes Medical Institute Janelia Research Campus, Ashburn, USA.

### Imaging Setup

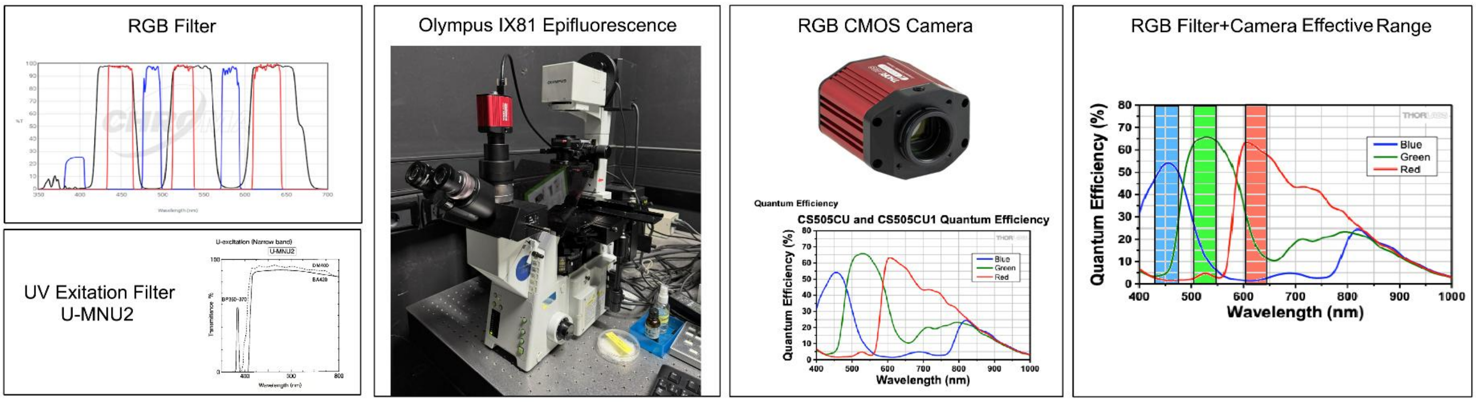


**Figure 1.S.** Imaging setup used in this work. Schematic representation of the optical configuration consisting of a white light source, optical filters, microscope, and RGB camera. Two alternative filter configurations were employed: (i) a multiband RGB filter cube (69015 ET – DAPI/Green/Red FISH; excitation 69015x, emission 69015m, dichroic 69015bs; Chroma Technology Corp.) for fluorescence excitation and detection in three bands (blue, green, red), and (ii) a modified UV-MNU2 filter used for autofluorescence imaging, where the emission filter was removed to collect the full emission spectrum. Images were acquired with an Olympus IX81 epifluorescence microscope equipped with a Thorlabs Kiralux CS505 5.0 MP RGB CMOS camera. The spectral response of the system results from the combination of the RGB filter transmission and the quantum efficiency of the camera detector, as shown in the

effective detection range plot. In the case of UV excitation with the modified filter, no emission filtering is applied; therefore, spectral selectivity is determined exclusively by the RGB camera response.

### Results

#### Clustering segmentation in the RGB Phasor Plot

##### Automatic segmentation using clustering techniques in the phasor space of RGB

This section provides a detailed discussion of the results presented throughout the paper, elaborating on the rationale behind the methods and algorithms chosen, particularly the clustering approaches. Specifically, for the phasor-based segmentation in hematoxylin and eosin (H&E) stained images, we applied several clustering algorithms to analyze their performance and measured the results presented in Table 1. These clustering algorithms were evaluated based on their segmentation efficiency using metrics such as Dice Similarity Coefficient, Jaccard Index, and Accuracy. These metrics allowed us to quantitatively compare the algorithms and assess their ability to accurately segment phasor plots in the context of H&E images. Clustering was chosen as it is an unsupervised learning approach, making it suitable for segmenting phasor plots without prior knowledge of labeled data. This is particularly useful in H&E images, where tissue features often exhibit complex spectral and spatial patterns. By leveraging clustering, we aimed to group similar spectral components into distinct clusters, facilitating the segmentation process. K-means was employed due to its simplicity and efficiency in segmenting data into a predefined number of clusters (k). It minimizes the variance within each cluster by iteratively adjusting cluster centroids, making it effective for datasets where clusters are roughly spherical and well-separated. Agglomerative Clustering is a hierarchical clustering method used because it constructs a tree-like structure (dendrogram) to represent nested clusters. It offers flexibility in capturing different cluster shapes and sizes. However, its sensitivity to the choice of linkage criteria (e.g., single, complete, or average) and initialization conditions were carefully considered during evaluation. Spectral Clustering was applied to exploit the local relationships between data points.

By constructing a similarity graph and using eigenvectors of its Laplacian matrix, this method can segment data with non-convex shapes and complex boundaries. Its ability to handle datasets with irregularly distributed clusters made it a valuable choice for phasor-based segmentation. GMM was chosen for its probabilistic nature, which models the data as a mixture of Gaussian distributions. This method allows clusters to have varying shapes and orientations by adjusting covariance structures (e.g., spherical, tied, or full), making it highly adaptable to the spectral diversity in H&E images.

For the data in Figure 1.B, we implemented the following clustering methods using Scikit-learn to segment the phasor plot in Figure 1.C: K-means with three components, Agglomerative Clustering with three components, Spectral Clustering with three components using nearest neighbor affinity, and Gaussian Mixture Model (GMM) with three different covariance types (Spherical, Tied, and Full). The segmentation results for K-means, Spectral Clustering, and GMM are shown in Figure 1.S. Each method produced an equally distributed phasor plot with three components, though there were slight phase shifts among them. As a result, the pseudocolor images derived from these methods exhibited the same phase shifting in the classified pixels. Consequently, the primary difference between these methods lies in the borders between components. There is no evidence to suggest a correct phase shift or starting phase value for each component; therefore, these methods can be used interchangeably to produce a uniformly distributed phasor plot. On the other hand, the Agglomerative method, being a hierarchical approach, is highly dependent on the initial conditions, which can lead to an inhomogeneous distribution in the phasor plot, as shown in the figure.

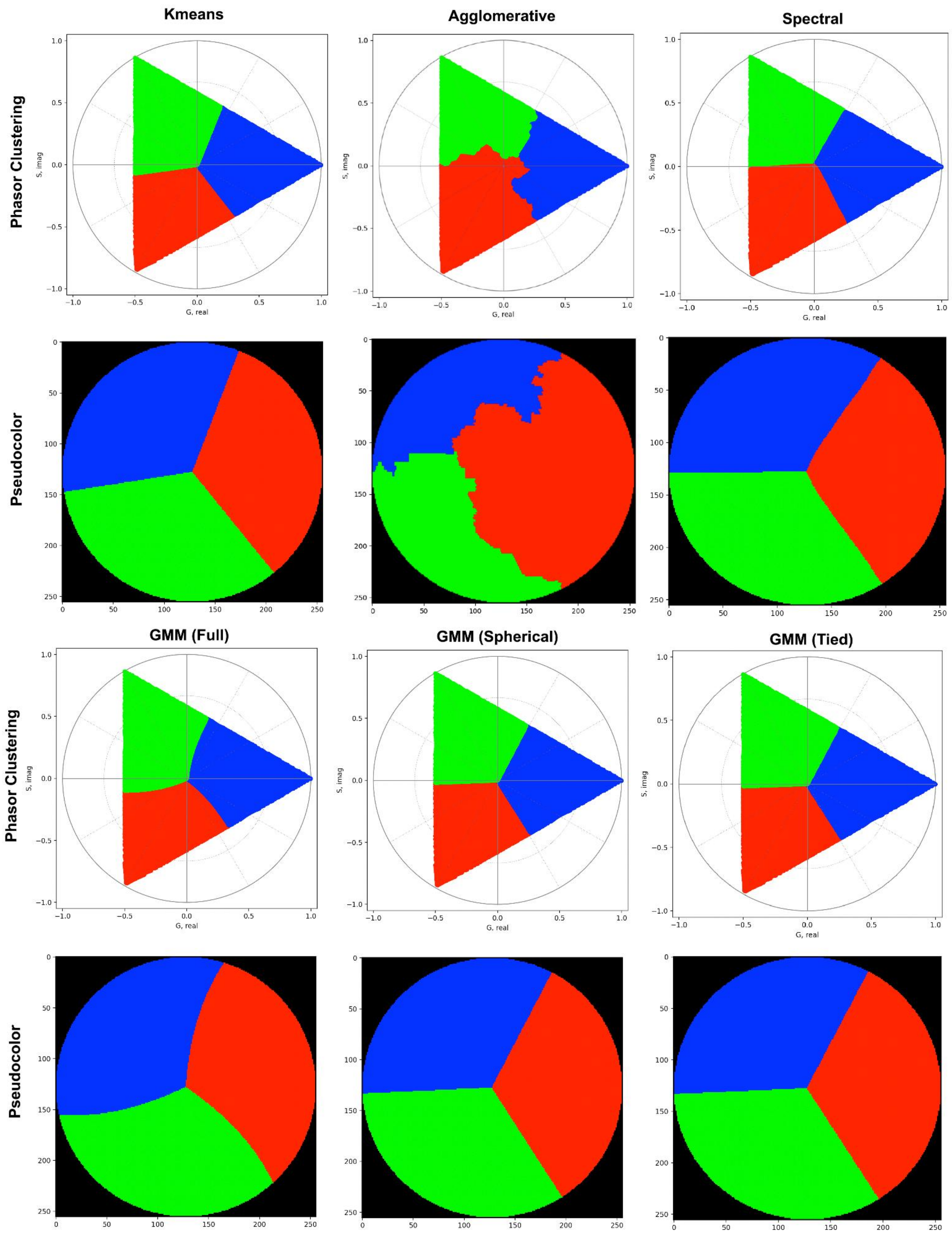


**Figure 2.S** Shows clusterized phasor plots obtained with different clustering methods and their corresponding pseudocolor images.

#### Execution time

We tested four clustering algorithms: K-means, Gaussian Mixture Model (GMM) with full covariance, GMM with tied covariance, and Spectral Clustering: on a hematoxylin and eosin (H&E) image with dimensions 2048x2448 pixels to evaluate their execution times for clustering tasks. The timing measurements included computing the clustering for 4 clusters, obtaining the cluster labels for each pixel, and creating the mask using the reshape method. The results, summarized in Table 1, also include estimated total times for processing 200 images. The results show that K-means is the fastest algorithm, with an execution time of 0.86 seconds per image and an estimated total time of 172 seconds for 200 images. Its speed makes it highly suitable for large-scale segmentation tasks, particularly when computational resources or time are constrained. In contrast, GMM with full covariance required 16.94 seconds per image and an estimated 3388 seconds (~56 minutes) for 200 images, reflecting the computational cost of estimating a full covariance matrix for each cluster. GMM with tied covariance, while faster at 9.64 seconds per image and 1928 seconds (~32 minutes) for 200 images, still lagged significantly behind K-means due to the inherent complexity of the GMM approach. Spectral Clustering, however, could not be computed for the given image. The failure to construct a fully connected similarity graph highlighted a key limitation of this method: it requires a well-connected graph to perform clustering. This issue underscores the need for preprocessing steps to ensure graph connectivity, which would further increase its computational overhead. It is important to note that the total estimated time for processing 200 images was calculated under the assumption that the computation time per image remains constant across iterations. However, this assumption may not hold true in practice. Variations in computational performance can occur as more images are processed, due to factors such as process accumulation, memory usage, or resource contention. These factors can lead to longer execution times for later iterations, particularly for methods like GMM that have higher memory and computational demands. As such, the estimated total times provide a useful baseline for comparison but should be interpreted with caution, as actual times may deviate depending on the system load and resource management. Overall, the findings emphasize the superior efficiency and scalability of K-means, which makes it the most practical choice for high-resolution images or large datasets. While GMM methods offer greater flexibility for modeling complex cluster shapes, their slower execution times limit scalability. Full covariance GMM, in particular, incurs a significant computational burden, while tied covariance GMM offers a trade-off with moderate gains in efficiency. Spectral Clustering's inability to execute highlights its sensitivity to data characteristics, such as graph connectivity, and suggests that its application requires additional consideration and preprocessing. In conclusion, K-means is the most efficient and practical algorithm for clustering tasks involving large datasets, especially when computational speed is critical. GMM methods, while slower, remain valuable for scenarios requiring more sophisticated modeling of cluster shapes. Meanwhile, Spectral Clustering presents challenges that must be addressed before it can be reliably applied, such as ensuring graph connectivity. These results provide clear guidance on the suitability of different clustering methods based on execution time and scalability for phasor-based segmentation tasks, while highlighting the need to account for real-world performance variability in estimating total computation times.

**Table 1.S.** Calculated and estimated execution time for each clustering algorithm for one image and for 200 images.

| | Kmeans | GMM (full) | GMM (tied) | Spectral |
|---|---|---|---|---|
| Time (1 image) | 0.86 s | 16.94 s | 9.64 s | Not computed |
| Total (estimated) | 172 s | 3388 s | 1928 s | - |

#### Methods comparison

The performance of the clustering methods was evaluated using three key metrics: Dice Similarity Coefficient, Jaccard Index, and Accuracy, as shown in Table 1. These metrics provided a quantitative comparison of segmentation efficiency between Multi-Otsu, K-Means, and Phasor-based methods for phasor-based segmentation in hematoxylin and eosin (H&E) stained images. By assessing overlap, union, and overall accuracy of the segmented regions, we determined how these methods performed relative to one another. The comparison between Multi-Otsu and K-Means revealed that K-Means achieves slightly better results across all metrics. Multi-Otsu vs. K-Means yielded a Dice score of 0.993 ± 0.003, a Jaccard score of 0.986 ± 0.005, and an Accuracy of 0.993 ± 0.002, indicating near-perfect segmentation with minimal variability. These results highlight K-Means' ability to deliver highly consistent, precise segmentation, making it particularly effective in scenarios with well-defined clusters. Multi-Otsu's strong performance in this comparison also underscores its reliability as a baseline method for phasor-based segmentation. When comparing Multi-Otsu to Phasor-based methods, results showed that while Phasor methods performed slightly lower in overall accuracy, they remained competitive and useful for segmentation tasks. Phasor-based methods achieved a Dice score of 0.951 ± 0.024, a Jaccard score of 0.907 ± 0.038, and an Accuracy of 0.953 ± 0.032. These values demonstrate that Phasor methods can effectively segment data, with a slight trade-off in precision and increased variability compared to Multi-Otsu. However, Phasor methods offer a distinct advantage: their reliance on phase and modulation information provides unique insights into spectral relationships that traditional intensity-based methods, such as Multi-Otsu, cannot capture. This makes Phasor methods especially valuable in applications where spectral characteristics are critical. The Phasor vs. K-Means comparison showed a similar trend. Phasor-based methods achieved a Dice score of 0.954 ± 0.024, a Jaccard score of 0.913 ± 0.039, and an Accuracy of 0.956 ± 0.032, slightly lower than K-Means but still within a competitive range. K-Means, with its centroid-based clustering approach, excelled in capturing well-separated clusters with high precision. However, Phasor methods offer flexibility in handling more complex spectral distributions, making them a useful alternative depending on the data characteristics.

The results show that K-Means consistently outperformed both the Multi-Otsu and Phasor-based methods in segmentation accuracy and precision, achieving the highest Dice, Jaccard, and Accuracy scores across all comparisons. This makes K-Means a highly reliable choice for segmenting phasor plots, particularly when the clusters are well-defined and relatively homogeneous. However, Phasor-based methods demonstrated strong performance, remaining competitive with Multi-Otsu and K-Means despite slightly lower scores. Their reliance on phase and modulation information provides unique advantages, especially when spectral data complexity or relationships are central to the analysis. While the higher variability of Phasor-based methods (\pm 0.024 to \pm 0.039) indicates sensitivity to data characteristics, their ability to adapt to more intricate spectral patterns makes them a valid and useful choice for specialized applications. Multi-Otsu, while reliable and consistent, may not capture the full spectral complexity that Phasor methods can address. However, its precision and ease of use make it a useful baseline for phasor-based segmentation tasks. In summary, while K-Means offers the best overall performance, Phasor-based methods are valuable alternatives that provide unique insights and flexibility, particularly when the analysis requires consideration of spectral relationships beyond intensity-based segmentation.

**Table 2.S.** Coefficient for each method compared 1-1.

| Method 1 | Method 2 | Dice | Jaccard | Accuracy |
|---|---|---|---|---|
| Multi Otsu | KMeans | 0.993 ± 0.003 | 0.986 ± 0.005 | 0.993 ± 0.002 |
| Multi Otsu | Phasor | 0.9510 ± 0.024 | 0.907 ± 0.038 | 0.953 ± 0.032 |
| Phasor | KMeans | 0.954 ± 0.024 | 0.913 ± 0.039 | 0.956 ± 0.032 |

**Areas obtained**

We analyzed the performance of three clustering methods: K-means, Multi-Otsu, and Phasor-based segmentation: on a dataset of 200 images corresponding to mouse lung alveolar regions. The dataset included two experimental groups: normal diet (ND) and normal diet instilled (ND inst), with 10 mice in each group and 10 images per mouse. The primary objective was to measure the areas of the alveolar regions segmented by each method and compare the results across the groups and methods. The findings, summarized as mean ± standard deviation, are as follows: for the ND group, K-means produced an area of 34.96 ± 5.93, Multi-Otsu yielded 34.81 ± 5.99, and Phasor-based segmentation resulted in 32.84 ± 5.78. For the ND inst group, K-means and Multi-Otsu yielded nearly identical areas of 69.18 ± 11.11 and 69.17 ± 11.19, respectively, while the Phasor-based segmentation estimated a slightly smaller area of 64.99 ± 11.23. For the ND group, K-means and Multi-Otsu showed highly consistent results, with mean areas differing by less than 0.2%. Phasor-based segmentation, while approximately 6% lower, still produced results within the range of variability observed across methods, demonstrating its reliability. A similar pattern was observed in the ND inst group, where K-means and Multi-Otsu again yielded almost identical area measurements, whereas Phasor-based segmentation provided a slightly more conservative estimate, approximately 7% lower. These differences may reflect the Phasor method's sensitivity to specific spectral features, which could influence its segmentation decisions and result in smaller detected regions. When comparing the ND and ND inst groups, all methods consistently captured a significant increase in alveolar areas in the ND inst group, with areas approximately doubling compared to the ND group. This consistency highlights the robustness of all three methods in detecting physiological or pathological changes in alveolar regions. The agreement between K-means and Multi-Otsu across both groups reinforces their suitability as reliable, interchangeable methods for segmenting these regions. Phasor-based segmentation, although slightly more conservative in its estimates, provides a valuable complementary approach due to its unique reliance on spectral data, which may reveal additional insights in specialized cases. In conclusion, K-means and Multi-Otsu are highly efficient and consistent methods for segmenting mouse lung alveolar regions, offering nearly identical results across all conditions. Phasor-based segmentation, while slightly underestimating areas, remains a valid and reliable alternative, particularly when spectral features play a critical role in the analysis. All three methods successfully detected significant differences between the ND and ND inst groups, making them suitable for identifying physiological changes in alveolar regions of the lung.

**Table 3.S.** Mean percentage of tissue areas obtained with each of the three methods apply.

| | **Kmeans** | **Multi Otsu** | **Phasor** |
|---|---|---|---|
| ND | 34.96 ± 5.93 | 34.81 ± 5.99 | 32.84 ± 5.78 |
| ND inst | 69.18 ± 11.11 | 69.17 ± 11.19 | 64.99 ± 11.23 |

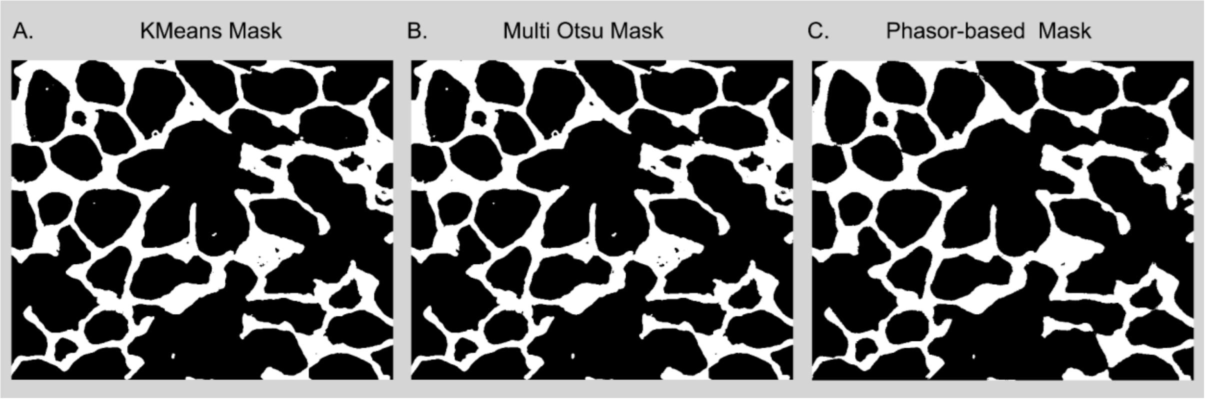


**Figure 3S.** Comparison of segmentation masks obtained with (A) K-means, (B) Multi-Otsu, and (C) phasor-based RGB analysis. While K-means and Multi-Otsu produce highly similar masks, the phasor-based method removes small isolated regions, yielding a cleaner segmentation with reduced pixel noise.

The comparison of segmentation approaches reveals that K-means (A) and Multi-Otsu (B) produce highly consistent masks, with only minor variations in boundary delineation. In contrast, the phasor-based segmentation (C) provides a cleaner representation by suppressing small isolated pixel clusters, effectively acting as a noise-filtering step. This results in smoother region boundaries and reduced fragmentation, while preserving the overall structural integrity of the segmented tissue.

### Autofluorescence Analysis

##### Using Principal Component Analysis

To quantify differences in spectral heterogeneity, we applied Principal Component Analysis (PCA) to the phasor coordinates of each condition. From the resulting PCA distribution, we computed the following descriptors:

- Variance 1 ($\sigma_1^2$): variance along the major axis of the PCA ellipse.
- Variance 2 ($\sigma_2^2$): variance along the minor axis.
- Elongation: defined as the ratio: $\frac{\sigma_1 2}{\sigma_2 2}$.
- Ellipse area: calculated as $A = \pi \sigma_1^2 \sigma_2^2$

These parameters were used to compare phasor spread and anisotropy across conditions and across acquisition methods (RGB vs HSI).

Spectral Phasor Entropy. Spectral complexity was assessed using a phasor-based entropy measure. The 2D phasor histogram (G-S plane) was normalized to form a probability distribution $\rho_i$ and the Shannon entropy was computed as:

$$H = \sum_i \rho_i \, log(\rho_i) \qquad (3)$$

This metric reflects the diversity of spectral components in the image: higher entropy indicates a broader range of spectral content and increased tissue heterogeneity.

Principal Component Analysis (PCA) reduces the phasor space into a simplified representation described by a few quantitative parameters, enabling direct comparison between imaging methods. PCA was applied to both melanoma and nevi datasets acquired in RGB and HSI. For melanoma, RGB images yielded an ellipse elongation of 21.8, with variances of 0.0304 and 0.0014 along the major and minor axes, respectively, and an ellipse area of 0.0204. In HSI, the elongation was 13.6, with variances of 0.0198 and 0.00145, and an ellipse area of 0.0168. For nevi, elongation values were lower, reaching 2.17 in RGB and 3.45 in HSI. The corresponding major axis variances were 0.00206 (RGB) and 0.00245 (HSI), while the minor axis variances were 0.00095 and 0.00071. The ellipse areas for nevi were 0.0043 (RGB) and 0.0042 (HSI). Entropy analysis of the phasor histograms further supported these findings, with melanoma showing higher entropy values (4.15 in RGB and 3.96 in HSI) compared to nevi (2.65 in RGB and 2.50 in HSI). PCA provided descriptors such as elongation, variance, and ellipse area, which consistently showed higher values in melanoma compared to nevi, reflecting greater spectral heterogeneity. These trends were observed in both RGB and HSI data, with moderate modality-dependent differences. Entropy analysis further supported these findings, with melanomas exhibiting higher phasor entropy than nevi in both modalities, indicating increased spectral variability. Together, these metrics provide objective evidence that melanoma lesions display enhanced spectral dispersion, reinforcing the qualitative differences observed in the phasor plots and pseudocolor maps.

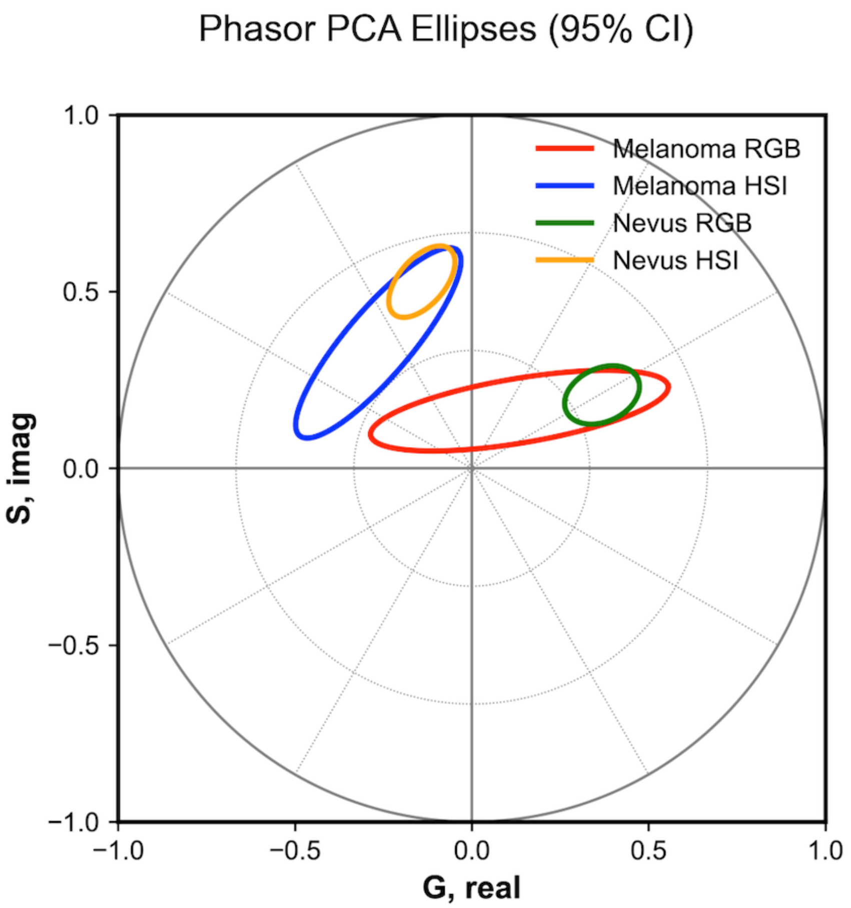


**Fig 4.S** Principal component analysis (PCA) applied to phasor distributions, showing 95% confidence ellipses for melanoma and nevus samples in both RGB and HSI modalities. Differences in shape, orientation, and dispersion reflect distinct spectral characteristics.

#### Spectral Unmixing

##### Components

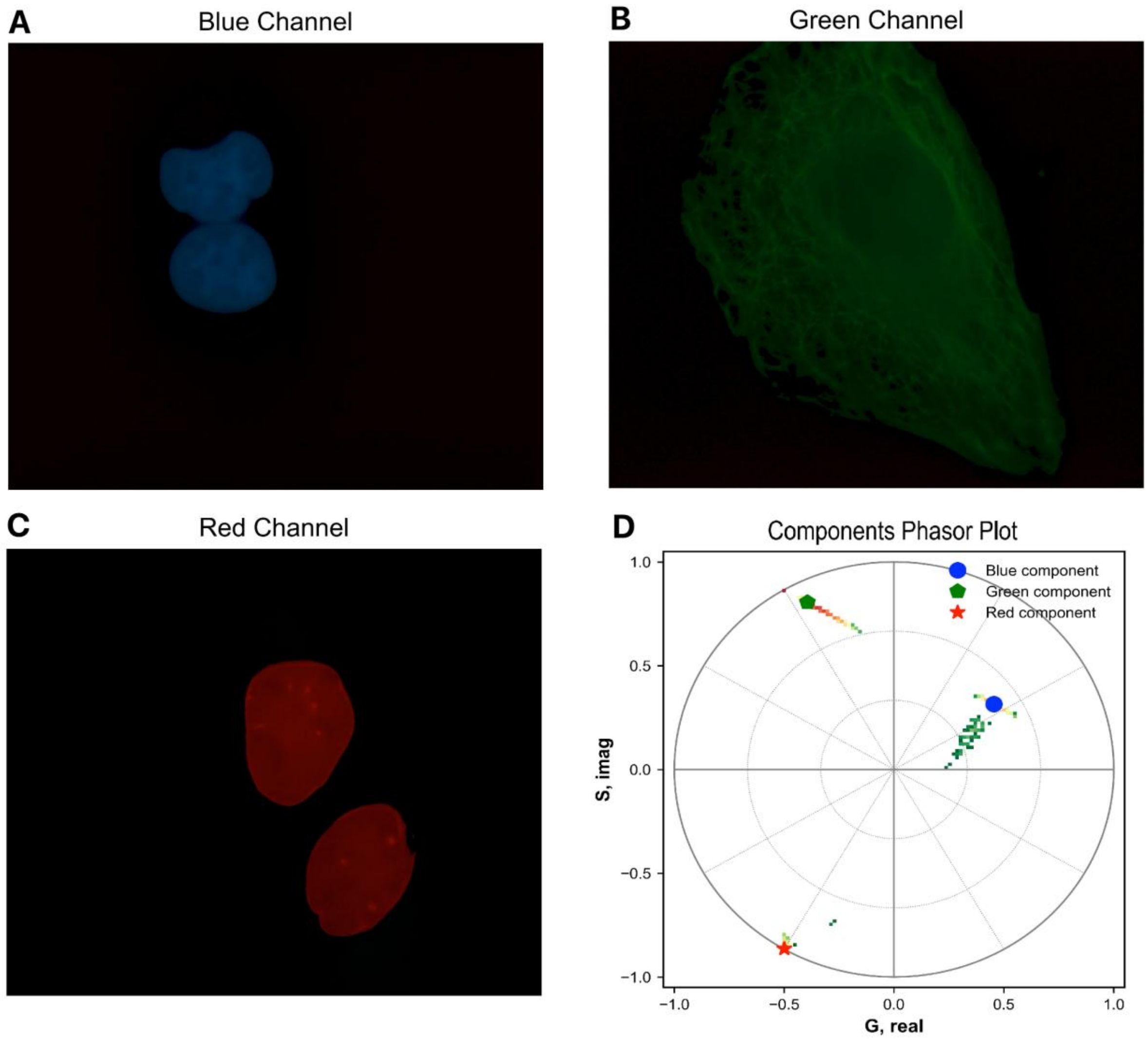


**Figure 5S. Phasor positions of pure fluorophores used for unmixing.**

(A–C) Individual images of pure-labeled samples: DAPI (blue) for nuclei, Tubulin-488 (green) for cytoskeleton, and Laminin-555 (red) for nuclear envelope. (D) Phasor plot of spectral signatures with their respective center of mass, used as anchor points for component separation in cell imaging, where the colored symbols mark the center of mass of the phasor distribution for each component: blue dot for DAPI, green hexagon for Tubulin-488, and red star for Laminin-555.

#### Intensity profile

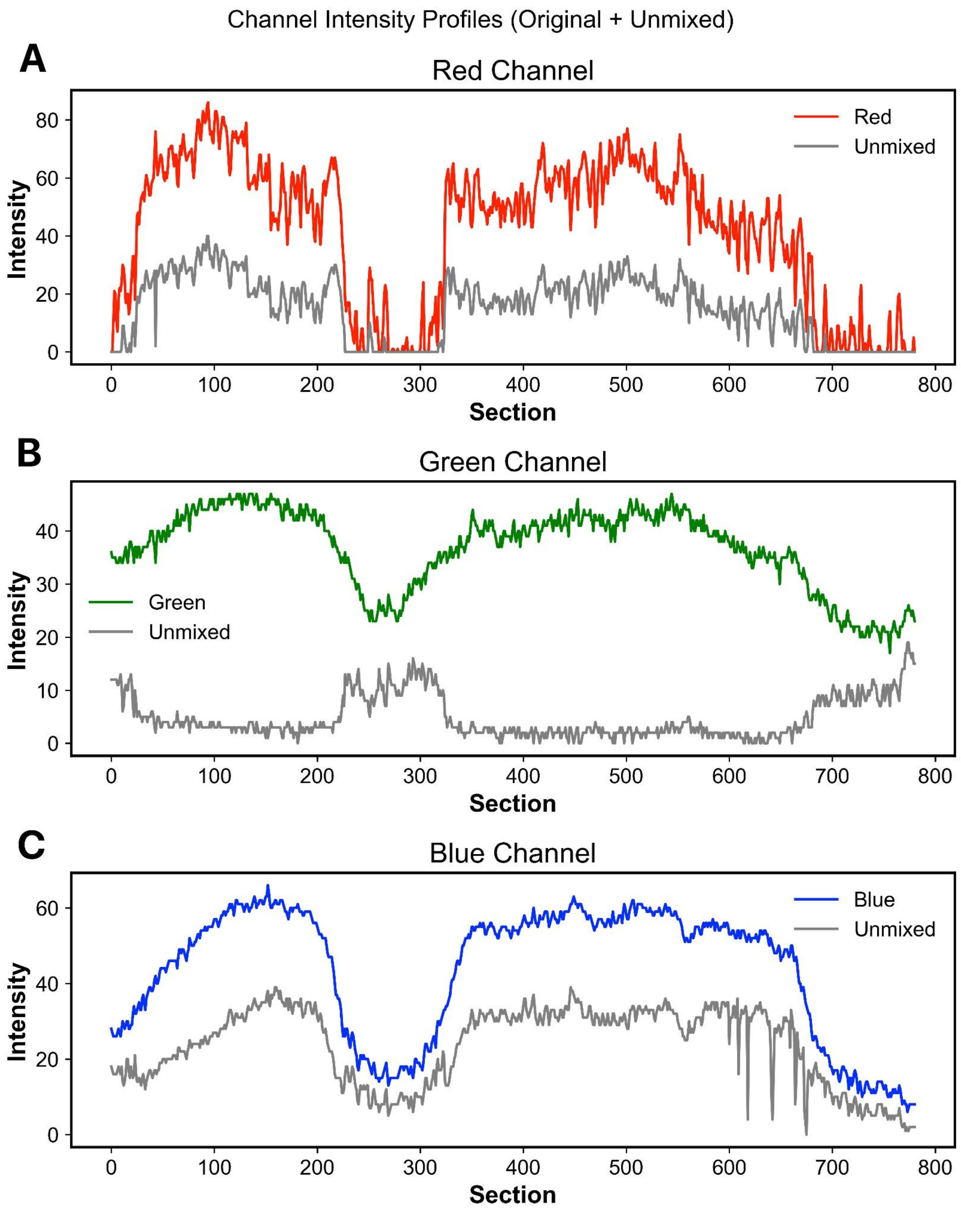


**Figure 6S. Intensity profiles of RGB channels in original and unmixed images.** (**A–C**) Intensity profiles of the red (A), green (B), and blue (C) channels measured along a selected line across the image. Colored lines represent the original RGB signal, while gray lines correspond to the unmixed signal obtained through phasor-based unmixing. The unmixed profiles reveal a smoother, component-resolved signal for each fluorophore, with reduced spectral crosstalk compared to the original channels.
